# Effective psychological treatment for PTSD changes the dynamics of specific large-scale brain networks

**DOI:** 10.1101/2020.01.07.891986

**Authors:** Marina Charquero-Ballester, Birgit Kleim, Diego Vidaurre, Christian Ruff, Eloise Stark, Jetro J. Tuulari, Hugh McManners, Yair Bar-Haim, Linda Bouquillon, Allison Moseley, Steven C. R Williams, Mark Woolrich, Morten L Kringelbach, Anke Ehlers

## Abstract

Very little is known about the role of effective cognitive therapy in reversing imbalances in brain activity after trauma. We hypothesised that exaggerated threat perception characteristic of post-traumatic stress disorder (PTSD), and subsequent recovery from this disorder, are underpinned by changes in the dynamics of large-scale brain networks. Here, we use a novel data-driven approach with high temporal precision to find recurring brain networks from fMRI data and estimate when these networks become active during exposure to either trauma reminders or neutral pictures. We found that PTSD patients spend less time in two default mode sub-networks in contrast to trauma-exposed healthy controls, and that PTSD symptom severity correlates positively with time spent in the salience network during exposure to trauma reminders. The former are important for different aspects of self-referential processing and the latter for detection of threat. Importantly, the decreased time in the default mode sub-networks is rebalanced after successful cognitive therapy for PTSD. Our results show that remittance of PTSD through trauma-focused cognitive therapy is associated with the successful reinstatement of a healthy balance in self-referential and threat detection brain networks.

## Introduction

Cognitive behaviour therapy (CBT) is one of the first-line treatments for a very wide range of mental health disorders (e.g., depression, posttraumatic stress disorder (PTSD) and anxiety disorders). Nowadays such conditions constitute one of the leading causes of disease burden worldwide (Vigo et al., 2016; Vos et al., 2015). Furthermore, it is estimated that overall treatment response rates averaged across anxiety disorders is approximately 50% at post-treatment (Loerinc et al., 2015), with about 20-40% of people still meeting criteria for PTSD or depression after CBT (Blanchard et al., 2003; Bradley et al., 2005; Holtforth et al., 2019). A mechanistic explanation of the effects of CBT in the brain, which is currently missing, is important to understand why this is the case, how to predict who will or will not respond to treatment, and how to help the group of individuals who do not respond.

This study investigated the effects of cognitive therapy for PTSD (CT-PTSD, (Ehlers et al., 2005)), one of the trauma-focused CBT programmes recommended by international treatment guidelines on the basis of its efficacy (e.g., American Psychological Association, 2017; International Society of Traumatic Stress Studies, 2018; National Institute for Health and Care Excellence, 2018). One of the hallmark symptoms of PTSD is the re-experiencing of the traumatic event (Brewin et al., 2010; Keane et al., 1997), that can be triggered by a wide range of trauma reminders and, very specifically, carries a feeling of “here and now” (Halligan et al., 2003; e.g., Michael et al., 2005, Bar-Haim, in prep). This is thought to indicate decontextualized retrieval of trauma memories (Brewin, 2016; Ehlers and Clark, 2000). The treatment used in this study builds on Ehlers and Clark’s (2000) cognitive model of PTSD and includes several specific procedures addressing the disjointed and decontextualised nature of trauma memories as well as problematic appraisals and coping strategies. It includes a specific discrimination training where patients learn to identify triggers of re-experiencing, and to focus on how the present trigger and its current context (“now”) is different from the trauma (‘then’). It further addresses excessively negative appraisals of the trauma and/or its sequelae, as well as of other people’s responses after the event, by providing information, Socratic questioning, updating the memories for the worst moments of the trauma by linking them with information that makes their meanings less threatening, and behavioural experiments. These strategies rely on the patient’s ability to mentalise, that is, to consider the thoughts, beliefs and feelings, both their own and those of other people, that underlie specific behaviours.

At the brain level, recent studies using functional magnetic resonance imaging (fMRI) suggest that three specific large-scale brain networks and their interactions play an essential role in both the healthy response to acute stress (Hermans et al., 2014, 2011; Oort et al., 2017) and the development of a wide range of psychiatric disorders, including PTSD, depression and generalised anxiety (Akiki et al., 2018, 2017; Menon, 2011). These three large-scale brain networks are the salience network (essential for cognitive processing of salient stimuli and interoception), the default mode network (DMN, attributed to self-referential processing) and the central executive network (involved in higher-order cognitive functions). In particular, acute stress is thought to lead to strategic reallocation of resources towards the salience network, shifting the configuration of these large-scale brain networks and explaining the behavioural changes observed in stressful situations (Hermans et al., 2011; Oort et al., 2017). Further, observed hypoactivation of the DMN in participants who go on to develop PTSD after a traumatic experience has been attributed to the destabilisation caused by a hyperactive or hyperconnected salience network (Akiki et al., 2017; Sripada et al., 2012a).

However, what exactly do we usually mean when we say that a network is hyperactive or hyperconnected? And which tools do we have to quantify changes in reallocation of resources to a particular network? Most PTSD fMRI studies looking at large-scale networks to date have used ‘time-averaged’ approaches, in which activation or connectivity (i.e., synchronisation among the regions within a network) is based on fMRI signal averaged over several minutes (e.g., Miller et al., 2017; Sripada et al., 2012a). As a result, any derived measures refer to a combination of the *total amount of time points* during which a network is active *(i.e., activity time)* or synchronised, and the *strength* with which this occurs each time. While this approach, by being so comprehensive, explains the largest proportion of variability in the fMRI signal between individuals, recent work suggests that it is the variance contained in the activity time of these networks that relates most strongly to affective traits such as fear or anger (Vidaurre et al., 2019). Furthermore, previous fMRI studies offer support in favour of a more constrained dynamic repertoire in PTSD participants in contrast to healthy controls (Mišić et al., 2016; Ou et al., 2014), which would presumably be reflected in the activity times of one or more large-scale brain networks. Additionally, by considering the evolution of dynamics across the entire scanning period during a particular experiment, we may explore the potential cumulative effects of a particular environmental exposure on brain activity.

Here we aim to identify the networks showing differences in their activity time(s) in PTSD participants in contrast to trauma-exposed controls without PTSD, and further hypothesise that symptom reduction through CT-PTSD will be accompanied by rebalancing of this particular temporal feature. This could potentially shed some light into the processes that lie at the core of PTSD and the specifics of how CBT operates on this disorder at the brain level.

To test our hypothesis, we apply a data-driven approach based on the use of Hidden Markov Models (HMM; Vidaurre et al., 2017b, 2016) on the fMRI data of a treatment-seeking PTSD population while exposed (1) to pictures of trauma reminders and (2) to neutral pictures. This approach allows us to estimate a (pre-specified) number of brain networks that recurrently activate across the scanning period – often referred to as brain states in the literature (e.g., Stevner et al., 2019; Vidaurre et al., 2018a, 2017b), and that are shared across our participants. Then, we calculate the overall amount of time during which each participant activates each brain network (i.e., activity time) in two different experimental conditions: trauma-related or neutral picture presentation. Together with fMRI data, information about the severity of their PTSD symptoms was acquired at each visit. For some participants visits occurred both before and after CT-PTSD (longitudinal sample) and, for others, only one single visit was completed, either before or after CT-PTSD. This added cross-sectional data to our sample, resulting in a mixed cross-sectional and longitudinal design. We refer to participants’ data acquired as part of their CT-PTSD at these two visits as pre-CT and post-CT. Some of the treatment-seeking participants were assigned to a waiting list, in which they were all tested at two points in time - 12 weeks apart on average - without having received any therapy in between (i.e., the pre-WAIT and post-WAIT data, serving as control for those collected pre-CT and post-CT). Additionally, same neuroimaging and behavioural data were acquired for a group of trauma-exposed but healthy participants (referred to as healthy controls) at a single point in time.

Supporting our hypothesis, we found that the activity time of two DMN sub-networks was increased up to levels similar to those of healthy controls after successful psychological treatment. Interestingly, only the activity time of one of the two DMN sub-networks – the medial temporal DMN, important for contextualised retrieval of autobiographical memories – appeared diminished in PTSD participants during presentation of trauma reminders and in relation to the severity of intrusive memories experienced during the fMRI scan. The activity time of the other DMN – the dorsomedial prefrontal DMN, important for mentalising –only showed significant correlations with PTSD symptomatology during presentation of neutral pictures. Further results revealed positive correlations between PTSD symptomatology and the activity times of salience and visual networks during presentation of trauma reminders.

## Results

### Successful cognitive therapy is associated with longer DMN activity

Analysis of the fMRI data using an HMM, which is an entirely data-driven approach, resulted in seven differentiated brain networks **(Figure 1a-1g)**. Each of these networks has a characteristic signature containing both spatial information (i.e., a specific combination of areas showing different degrees of activation relative to the average (+/-)) and temporal information (i.e., a networks time-course showing when these patterns become active). For two of these networks, the spatial map showed areas of above-average activation (+) consistent with the DMN. More specifically, one of the spatial maps was consistent with the medial temporal DMN (mtDMN^+^) **(Figure 1a)**, and the other one was consistent with the dorsomedial prefrontal DMN (dmPFC DMN^+^) **(Figure 1b)**. The other networks also resemble well-established functional networks. Details of the assessment of the states’ neuroanatomy are described in the methods and in **Supplementary figures 1 and 2**. An overview of the group average activity times for each network, separated for trauma-related and neutral picture blocks, is presented in **Supplementary Figure 3.**

**Figure 1.**
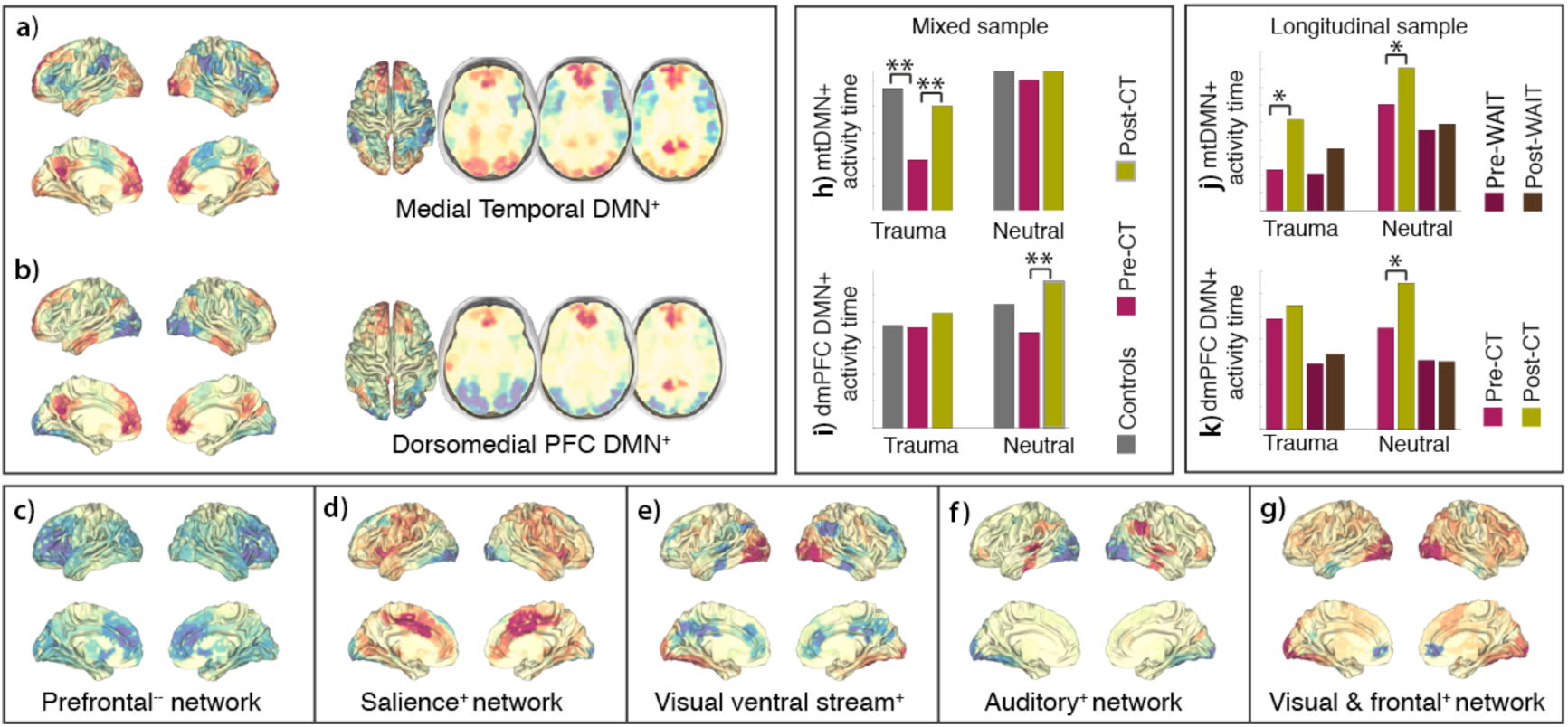
PTSD participants pre-therapy spend significantly less time activating DMN-related states in comparison with post-therapy and healthy controls. a-g) Spatial maps for our seven networks, labelled according to the areas of above (+) or below (-) average activation. h) During presentation of trauma-related pictures, PTSD participants pre-CT spend significantly less time activating the medial temporal DMN+ (mtDMN^+)^ than participants post-CT and healthy controls. i) During presentation of neutral pictures, PTSD participants spend significantly less time activating the dorsomedial PFC DMN^+^ (dmPFC DMN) before than after CT-PTSD. j-k) Increases in the time engaging the mtDMN^+^ and the dmPFC DMN^+^ were observable pre- and post-CT but not pre- & post-waiting list. c-g) No significant differences between groups were observed in the activation time of any of the other networks. *p < 0.05, **FDR < 0.05 corrected across group comparisons, experimental condition and number of networks.

This analysis included data from the mixed sample; that is, from participants that completed both sessions (before and after CT-PTSD) as well as participants that completed only one session (before or after CT-PTSD). The results presented were corrected across group comparisons, experimental conditions and number of networks. Results show that during the presentation of trauma reminders, pre-CT participants spent less time activating the mtDMN^+^ than post-CT (p=0.0126). We also observed a significant difference in activity time of the mtDMN^+^ between pre-CT and healthy controls (p=0.0168) in this condition, but no significant differences were observed between healthy controls and post-CT (p=0.7790) **(Figure 1h)**. This suggests that the therapy was successful in bringing activity time of the mtDMN^+^ back to the levels of healthy controls in the post-CT group when exposed to reminders of the traumatic event. No significant differences between groups were observed in activity time of the mtDMN^+^ during presentation of neutral pictures (**Figure 1h**).

However, during presentation of neutral pictures, we observed that pre-CT participants spent less time activating the dmPFC DMN^+^ than post-CT participants (p=0.0294) **(Figure 1i)**. No significant differences were observed between healthy controls and any of the PTSD groups, neither pre-CT nor post-CT. Also, no significant differences were observed between pre- and post-CT participants in the activity time of the dmPFC DMN^+^ during the presentation of trauma reminders **(Figure 1i).**

Hence, activity times of both the mtDMN^+^ and the dmPFC DMN^+^ differ between groups, but these differences are specific to the experimental condition. Differences in activity time of the mtDMN^+^ are specific to the blocks during which trauma reminders are presented, while differences in activity time of the dmPFC DMN^+^ appeared only during presentation of neutral pictures.

### Changes observed in activity time of the DMN after CT were not present in the waiting list group

We then ran a second analysis to investigate whether the increased activity time of the DMN networks observed in participants after CT was present in the group of participants that were assigned to a waiting list condition **(Figure 1j-1k).** This was a strictly longitudinal analysis, including only participants that had completed both scans: a first scan at baseline, followed by a second scan either after CT-PTSD (n=14) or after an average of 3 months on the waiting list (n=8). Results presented here are corrected across group comparisons, experimental conditions and number of networks; but in those cases in which significance was borderline, we also show uncorrected results. No significant changes in activity time of the mtDMN^+^ were observed when comparing the scan obtained at baseline to the scan obtained after 3 months of waiting, neither during presentation of trauma-related (p=0.3629), nor during presentation of neutral pictures (p=0.40). Findings of increased activity time of the mtDMN^+^ in the post-CT during presentation of trauma-related pictures were corroborated, although they were only borderline significant after correction for multiple comparisons (p=0.0525 after correction, p=0.008 uncorrected) (**Figure 1j**).

In relation to activity time of the dmPFC DMN^+^ during presentation of neutral pictures, we observed that the waiting list group did not show any differences across sessions (p=0.48), and the CT group showed only a non-significant increase in the time visiting the dmPFC DMN^+^ after CT once we had corrected for multiple comparisons (p=0.0952 after correction, p=0.0136 uncorrected) (**Figure 1k**).

No significant changes between the first and the second visit were found in the activity time of any of the other networks, neither after CT nor after waiting list.

To test for a possible interaction between treatment condition and time, we calculated the ‘amount of change’ in activity time for the mtDMN^+^ and the dmPFC DMN^+^ between the two visits, both for the pre-CT vs. post-CT and pre-Wait vs. post-Wait conditions. We then compared these values to assess whether there were any significant differences in the amount of change in the activity time of the two DMN networks linked to the treatment condition (CT-PTSD vs. Waiting list). This analysis yielded no significant results.

### Activity time of DMN and salience networks correlate with PTSD severity

To further investigate the association between network activity times and PTSD symptomatology, we computed correlations between the activity times of each of the networks and the severity of each of the three PTSD symptom clusters (i.e., Re-experiencing, Avoidance and Hyperarousal) as described in the DSM-IV (American Psychiatric Association, 2000). This was done in order to investigate whether time spent on any of the networks was related to any specific symptom clusters. We integrated different statistical tests into a single p-value using the non-parametric combination (NPC) algorithm (Vidaurre et al., 2018c; Winkler et al., 2016), which integrates all hypotheses into one, enabling us to test whether at least one alternative hypothesis is true. In particular, we aggregated p-values across symptom clusters, instead of across networks, in order to assess whether at least one of these symptom clusters bears a relationship to the activity time of any of the networks. Additionally, we also considered individual correlations corrected across multiple comparisons. As with previous analyses, we did this separately for the neutral and the trauma-related conditions. The analysis included participants from the therapy (pre- and post-CT) and the waiting-list groups.

We found that the activity times of the mtDMN^+^ and the salience^+^ networks were related to at least one of the symptoms clusters (p=0.006 for mtDMN^+^ and p=0.0399 for the salience^+^ network, respectively) during presentation of trauma-related pictures (**Figure 2a**). Also, the activity time of the dmPFC DMN^+^ was associated with at least one of the symptoms clusters (p=0.0046) **(Figure 2b)** during presentation of neutral pictures. Furthermore, closer examination revealed that each of the symptom clusters significantly correlated with the activity times of the mtDMN+ (*re-experiencing (p<0.002), avoidance (p<0.008) and hyperarousal (p<0.002)* **(Figure 2, column 1)**, dmPFC DMN+ (*re-experiencing (p<0.027), avoidance (p<0.045) and hyperarousal (p<0.004)* **(Figure 2, column 3**), and salience+ networks (*re-experiencing (p<0.033), avoidance (p<0.039) and hyperarousal (p<0.022)* **(Figure 2, column 2)**, after correcting across number of networks and number of symptom clusters.

**Figure 2.**
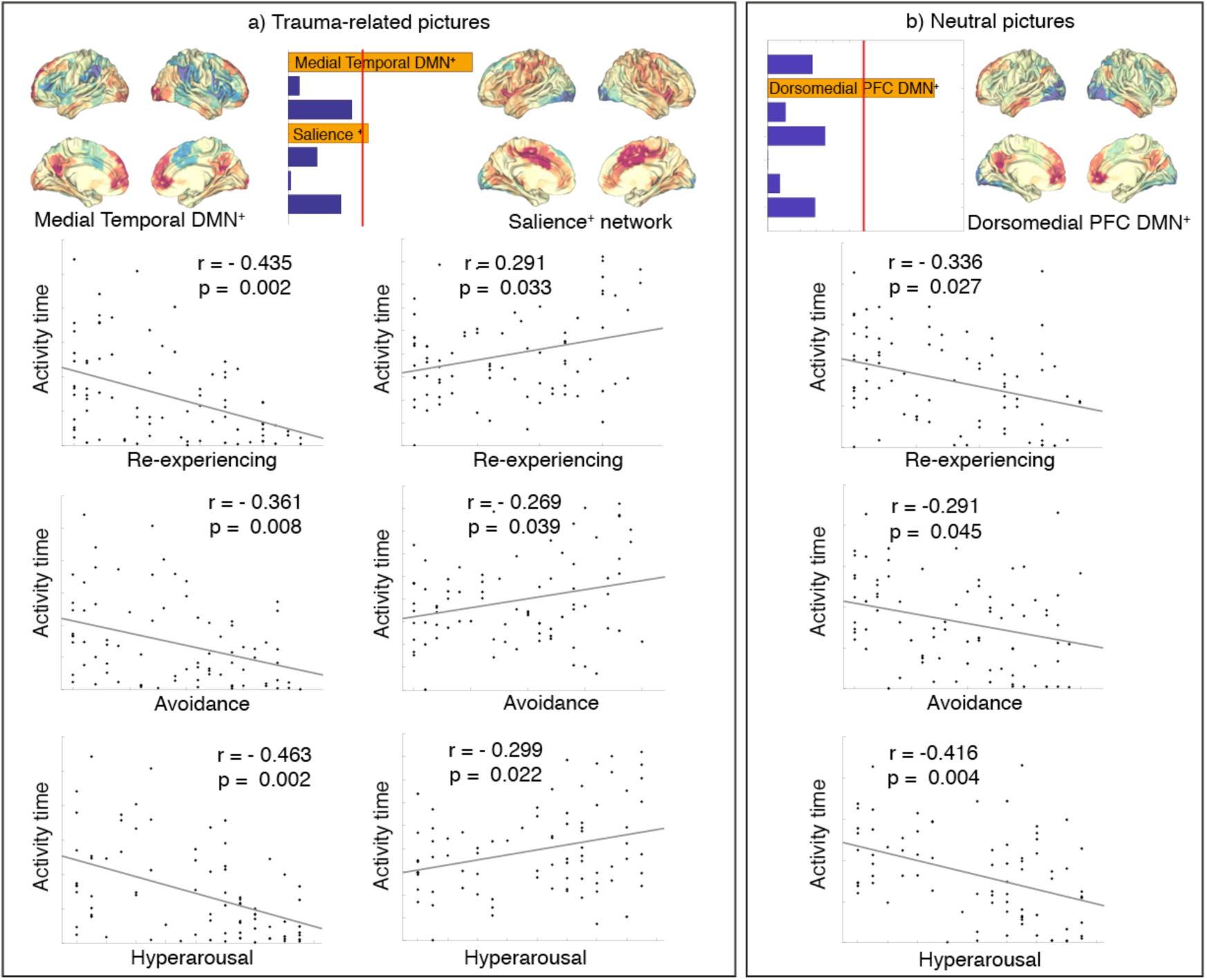
Symptom severity shows significant correlations with activity time of the sub-networks of the DMN, the salience network and each of the three symptom clusters (re-experiencing, avoidance and hyperarousal). a) Integration of different statistical tests into a single p-value through non-parametric combination algorithm reveals a significant relationship between PTSD symptoms and the activity times of the medial temporal DMN^+^ and the salience^+^ networks. More in detail, the first column shows the significant negative correlation between the activity time of the medial temporal DMN (mtDMN^+^) and re-experiencing (p<0.002), avoidance (p<0.008) and hyperarousal (p<0.002), while the second column shows the positive correlation with the activity time of the salience^+^ state and each of the symptom clusters: re-experiencing (p<0.033), avoidance (p<0.039) and hyperarousal (p<0.022). b) Statistical integration into a single p-value shows a significant relationship between PTSD symptoms and the activity times of the dorsomedial PFC DMN^+^. The third column shows the significant negative correlation between the activity time of the dorsomedial PFC DMN (dmPFC DMN^+^) and re-experiencing (p<0.027), avoidance (p<0.045) and hyperarousal (p<0.004).

Cluster-specific positive correlations during presentation of trauma-related pictures were found between the activity time of the prefrontal^-^ network and re-experiencing symptoms (p = 0.024) as well as between the visual/frontal^+^ and hyperarousal (p = 0.03). Activity time from the other brain networks did not show any significant relationship with PTSD symptomatology after correcting across number of networks and number of symptom clusters.

### The flashback-like qualities of intrusive memories in the scanner correlate with activity time of several brain networks

We explored the relationship between activity times and the flashback qualities (i.e., vividness, distress, and feeling of ‘here & now’) associated with intrusive memories characteristic of PTSD. When intrusive memories arise, the re-experiencing of the traumatic event occurs in a very vivid way, mostly consisting of visual impressions, and carries a feeling of the event occurring in the present moment with the consequent distress (Ehlers and Clark, 2000; Michael et al., 2005).

Analyses were carried out on the subsample of participants who experienced at least one intrusive memory during the fMRI task (n=72). Again we considered aggregated statistical tests into a single p-value through the NPC algorithm (Vidaurre et al., 2018c; Winkler et al., 2016), as well as individual correlations corrected across multiple comparisons. Results from correlations aggregated across flashback-like qualities revealed that the activity times of mtDMN^+^ and the salience network hold linear relationships with at least one of the flashback-like qualities **(Figure 3c, left)**. Individual correlations between the activity time of each of the networks and each of the flashback-like qualities revealed a negative correlation between mtDMN^+^ and memory-related distress (p=0.036 corrected) and a positive correlation between the salience^+^ and the feeling of ‘here & now’ (p=0.0607 after correction, p=0.0061 uncorrected). These were the only significant correlations after correction for multiple comparisons between the mtDMN^+^ and the salience networks, and specific flashback-like qualities. However, other qualities were significant before correction for multiple comparisons (i.e., vividness and the feeling of ‘here & now’ for the mtDMN^+^ and all other three variables for the salience network). Furthermore, we observed a positive correlation between the activity time of the ventral visual stream^+^ and distress, which was significant after correction for multiple comparisons **(Figure 3d)**.

**Figure 3.**
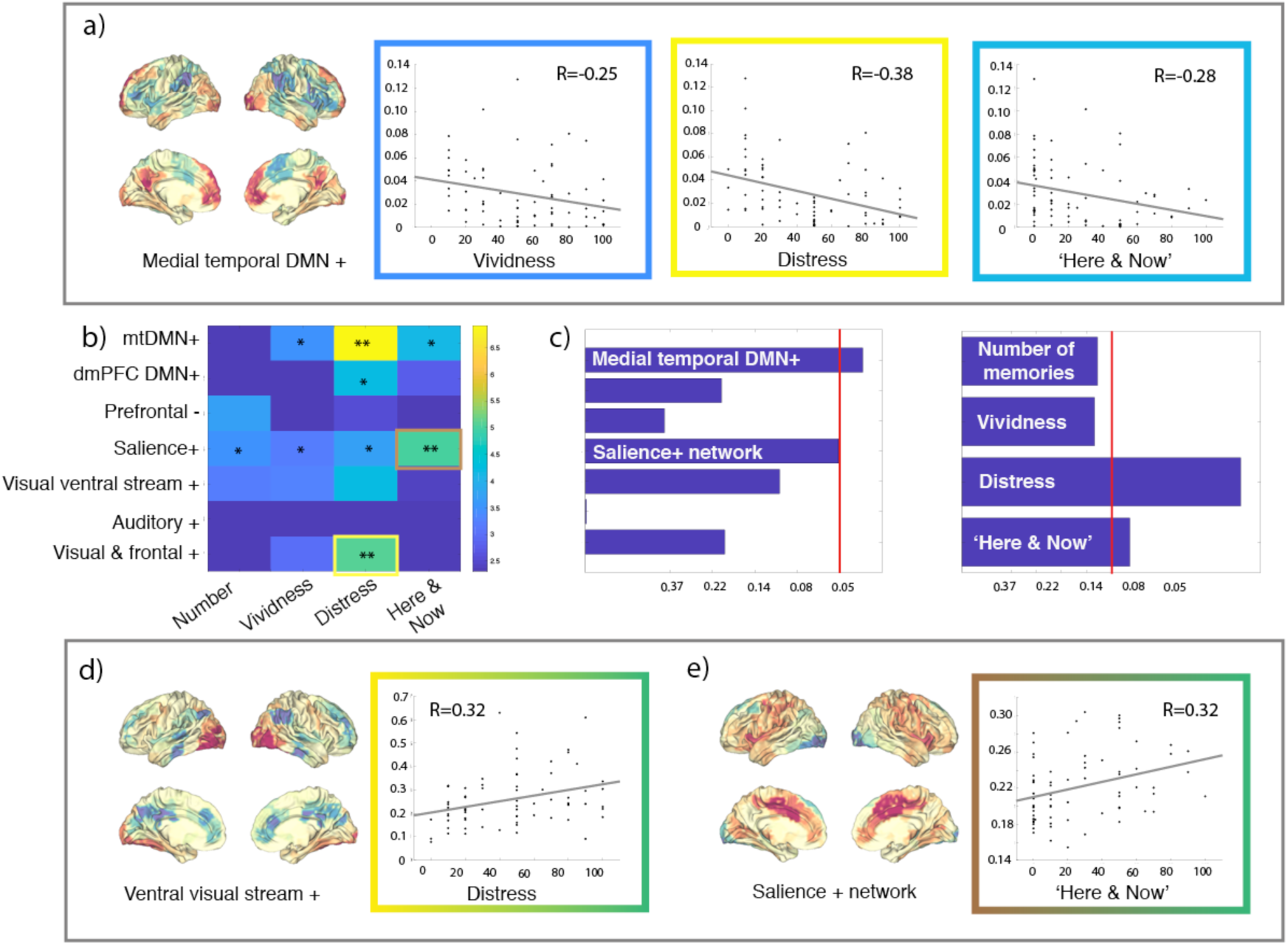
The activity time of different states is significantly correlated with the flashback-like qualities of intrusive memories in the scanner. a) Correlation between time spent in the ‘Medial temporal DMN^+^’, and quality of the intrusive memories. All variables related to quality of the memory are negatively correlated to time spent on this network, although only the relationship with how distressing the intrusive memories were remains significant after FWER correction. b) Matrix of correlation p-values for each of the network activity times with each of the memory variables (*p < 0.05 **FWER < 0.05). c) Aggregated p-values for the correlations of networks’ activity times across memory variables (left) and memory variables across networks’ activity times (right), FWER corrected. Time spent in the Visual ventral stream^+^ (d) and Salience^+^ networks (e) correlates with how distressing and how much the memory felt as if it was currently happening, respectively.

Further, we computed aggregated p-values across the seven networks for each of the flashback-like memory variables. Here, we observed that the degree of distress caused by the memory, and how much it made the participant feel as if it was happening again, could be predicted by a combination of the activity times of the estimated networks **(Figure 3c, right)**.

### Differences in activity time of the mtDMN^+^ might be due to amplification of the activity time in the mid and late stages of the task

We computed the average activity across participants for each point in time along the fMRI scan to see whether activity times might be influenced by long-term dependencies, i.e., by events prior to the one directly preceding the present point in time. From a cognitive perspective, such an effect could be reflecting a cumulative effect of exposure to trauma reminders on the activity time of specific networks.

First, we observed that most networks’ probabilities of activity time follow an oscillatory pattern with 40 cycles, corresponding to the 40 blocks of pictures that form the experimental task **(Supplementary figure 4)**. The only exception to this was the prefrontal^-^ network, where we observed short-lived ‘spikes’ of activity, perhaps related to sharp increases in the probability of this network becoming active in association with specific pictures within the picture blocks.

Second, it appears that the amplitude of the oscillations remains stable across the entire scan for most networks, except for the mtDMN^+^. Here, the probability of activity seems to increase after approximately the first 50 repetition times. This could indicate that the experimental task has a cumulative effect that reflects exclusively in the activity time of the mtDMN+.

Third, we observed that the oscillatory patterns show a high degree of overlap between groups (i.e., healthy controls, pre- and post-CT-PTSD) for most networks, but less so for the mtDMN^+^. These differences in the group-averaged activity time probability are coherent with the significant differences observed when comparing healthy controls, pre- and post CT-PTSD. Future work should assess, whether the differences between groups interact with time along the experimental task, and if controls and remitted PTSD show a stronger increase in the activity times of the mtDMN+ as the task progresses than in participants with current PTSD.

## Discussion

This study shows that successful CT-PTSD is associated with changes in the time spent in specific large-scale brain networks. We found that participants with PTSD spent significantly less time engaging two different DMN-related networks than did healthy participants: one network with a stronger temporal component (mtDMN^+^) and one with a stronger dorsomedial prefrontal component (dmPFC DMN^+^). Furthermore, the time the two networks were active in PTSD was increased after successful CT to levels similar to those of healthy participants. In line with these results, we found a significant negative correlation between severity of PTSD symptoms and time activating these DMN-related networks. This resonates with previous studies of DMN hypoactivity and hypoconnectivity in PTSD (Bluhm et al., 2009; King et al., 2016; Lanius et al., 2010; Sripada et al., 2012b). Our results further add to this literature by suggesting that activity time may be one of the network behaviours playing a role in the observed dysfunction in the DMN, and that this aspect was modified after successful CT-PTSD.

The data-driven estimation of two DMN networks is consistent with empirical papers and related theories that consider the DMN a constellation of several subsystems associated with different forms of self-referential processing (Andrews-Hanna, 2012; Andrews-Hanna et al., 2010; Jessica R. Andrews-Hanna et al., 2014; Archer et al., 2015; Northoff and Bermpohl, 2004; Vidaurre et al., 2018b). Beyond normalisation of the two DMNs activity time after CT-PTSD, our study shows spatial specificity within the DMN in relation to the experimental condition (trauma vs. neutral picture presentation) and the severity of intrusive memories in the scanner. In particular, we show that the strength of intrusive memories with flashback-like qualities, the hallmark symptom of PTSD, was only related to the activity time of the mtDMN^+^ during presentation of trauma-related pictures, but not of the dmPFC DMN^+^ **(Figure 3a)**. From the two networks, only the spatial map of the mtDMN^+^ showed above-average activation in parahippocampal cortex, hippocampal formation and visual cortex. Together with the midline structures (i.e., medial prefrontal cortex and precuneus), whose above-average activation was part of both DMN networks, the hippocampal and parahippocampal formations form a network that has been related to the contextualised retrieval of episodic memories (Andrews-Hanna et al., 2010; Jessica R Andrews-Hanna et al., 2014; Cabeza and St Jacques, 2007; Ranganath and Ritchey, 2012; Rugg and Vilberg, 2013).

In the context of the cognitive theory of PTSD (Ehlers and Clark, 2000), one of two factors resulting in the sense of current threat that perpetuates PTSD symptomatology is the spatially and temporally decontextualised memory of the traumatic event. The difficulty that PTSD participants have -in contrast to healthy trauma-exposed participants – in entering this network for a sufficient amount of time during presentation of trauma reminders, might be related to a decontextualised retrieval of traumatic memories. A previous study by St Jacques and colleagues (2013) on trauma-naïve and PTSD participants showed that participants who experienced memory retrieval of the traumatic event from the perspective of their own eyes (as opposed to experiencing it in a detached way from oneself) recruited the medial temporal lobe network more.

From a neuroscientific point of view, one could argue that deficient encoding of the traumatic event might have prevented migration of the memory from the hippocampal formation to a distributed network of brain regions during the consolidation process (Mcclelland et al., 1995; Squire and Alvarez, 1995). This ‘earlier’ form of the memory remains under-contextualised and, therefore, indiscriminately accessible; that is, an intrusive memory with flashback-like qualities. One of the goals of CT-PTSD is the development of a coherent narrative that places the series of events in chronological context and includes the more up-to-date information that the patient has now (e.g. ‘I did not die’ or ‘It was not my fault’). This is thought to reduce re-experiencing through the creation of an updated, contextualised memory that inhibits the cue-driven retrieval of intrusive memoires, and is experienced as an event in the past rather than the present (Conway, 1997; Ehlers and Clark, 2000). CT-PTSD focuses specifically on the worst moments of the trauma, which are represented in re-experiencing symptoms, and targets those parts of the trauma memory through a process of contextualising and updating. The result of this process is a contextualised memory, which might underlie the observed increase in activity time of the medial temporal DMN during presentation of trauma reminders after CT-PTSD.

Continuing with the cognitive theory of PTSD, the other main factor contributing to the sense of current threat is the presence of negative appraisals of the traumatic event (e.g., ‘What happened was my fault’ or ‘My family despises me because of what I did/didn’t do) (Ehlers and Clark, 2000). CT-PTSD identifies those appraisals and changes them through a variety of techniques that use our mentalisation processes (Björgvinsson and Hart, 2008); that is, our ability to understand the mental states that underlie our own and others overt behaviour. Specifically, the dmPFC DMN^+^ is considered to support mentalisation (Andrews-Hanna et al., 2010; Jessica R Andrews-Hanna et al., 2014). And so the observed increase in activity time of this network after CT-PTSD might be related to the work done during the psychological therapy.

A preliminary exploration of the activation patterns of mtDMN^+^ across participants further suggests a more marked progressive increase in activity time across the scan in healthy controls and remitted participants than in those with current PTSD **(Supplementary figure 4)**. Given the role of the mtDMN^+^ discussed in previous sections, we speculate that the progressive activity time increase of this network in healthy and recovered participants might be related to an increasing number of details being recalled along with the recurring exposure to trauma reminders.

Beyond the disruption of the DMN, previous neuroimaging literature links hyperactivity and hyperconnectivity of the salience network to PTSD (Abdallah et al., 2019; Sripada et al., 2012b). Our results are in line with these findings, as we show positive correlations between activity time of a salience network and severity of PTSD symptomatology. We show this in terms of the DSM-IV symptom clusters **(Figure 2)**, and also flashback qualities of intrusive memories **(Figure 3)** during the presentation of trauma-related pictures. Other networks with a strong visual component also show positive relationships between their activity time during presentation of trauma-related pictures and symptom severity. Overall, we interpret this as a result of increased saliency of trauma-related images for people with PTSD, accompanied by an increase in alertness that would lead to more frequent activity of perception and salience networks.

By contrast to studies that average the brain signal across the temporal dimension, the very much higher temporal precision we have used in this study enables exploration of theoretical questions regarding the stress response and psychiatric disorders. Recent theories based on animal research propose that the stress response entails dynamic shifts in network balance, with the DMN, the salience, and the central executive networks playing the key roles (Hermans et al., 2014, 2011; Oort et al., 2017). It has been further argued that some affective disorders such as PTSD are likely linked to disruptions in the shifts of these large-scale network (Akiki et al., 2017; Menon, 2011), which could manifest in an inability to disengage a specific pattern of brain activity and transition into another one. More recent work has shown that the variance contained in the fluctuations of these networks are more strongly related to affective traits such as fear or anger than the individual differences found in connectivity measures derived from the totality of the fMRI signal averaged over several minutes (Vidaurre et al., 2019). While much has been hypothesised about the dynamic behaviour of large-scale brain networks, appropriate tools to address these hypotheses directly in human brain data were underdeveloped. The data-driven approach used here, by allowing for concurrent estimation of both large-scale brain networks and the specific time periods during which these dominate brain activity, was able to provide evidence in favour of difficulties in engaging DMN components in participants with PTSD.

PTSD is a useful model for studying the neural changes that occur in the brain following CBT. While the focus of this work is on PTSD, it is important to note that PTSD shares many commonalities with other psychiatric disorders, such as depression or anxiety disorders (Brady et al., 2000; Gros et al., 2012). These disorders can also develop after a traumatic experience, are highly comorbid with PTSD, and are often treated through a range of CBT programmes (National Institute for Health and Care Excellence, 2011, 2009). Furthermore, theories linking affective disorders to disruptions in the dynamic shift of large-scale brain networks apply to these disorders as well (Menon, 2011). Here we explored the brain changes pre- and post-CT in a population of participants with PTSD, some of whom have comorbid depression and anxiety disorders. Our results, therefore, might be relevant to overarching processes important for regaining mental health through CT, going beyond the specifics of PTSD psychopathology.

In order to continue with the characterisation of large-scale network dynamics in health and disease, more studies using time-varying approaches are needed. These should consider some limitations of the present work. First, a larger longitudinal sample would be desirable in order to reach more robust conclusions related to the changes in activity time of the networks in the pre- and post CT and pre- and post- waiting list conditions. Second, the time spent engaging the mtDMN^+^ during presentation of trauma-related pictures in the mixed analysis **(Figure 1h)** was partly explained by a combination of differences in gender and medication when comparing healthy controls to pre-CT participants. However, this was not the case for pre-CT vs. post-CT, where the differences persisted even after controlling for these two variables. While in an ideal scenario all groups should be matched for all potentially confounding variables, this is not always possible – future studies should still aim to reduce these differences. Third, while we decided to focus on the trauma-related and neutral blocks of pictures presented, this is a simplification of our experimental task, which further included the presentation of 0-2 mushroom pictures per block. However, the present work did not consider the potential influence of the mushroom picture presentation in the activity times of the networks; this remains to be explored in future work. Fourth, our interpretation of the data-driven large-scale brain networks was supported by an automatic meta-analysis carried out through Neurosynth (Yarkoni et al., 2011) **(Supplementary figures 1 and 2)**. While we think that this has the potential to provide useful insights and to foster interesting questions for further research, we are also aware of the limitations of reverse inference based on literature mining. Last, we think that the field could greatly benefit from studies assessing the activity times of large-scale brain networks both in treatment responders and non-responders.

Overall this study shows that time-varying approaches may allow us to test some core aspects of cognitive and biological models of psychiatric disorders. Future studies could address the specifics of direct interactions between large-scale brain networks, in particular the salience and the DMN, to characterise further the dynamics related to PTSD symptomatology. Studies including treatment-resistant cases in their sample might be able to clarify how their dynamics differ from those who respond to treatment, and whether this is most strongly related to the activity time or the activation strength of specific large-scale brain networks. In due course, this could lead to a precise understanding of suboptimal mechanisms in treatment resistant cases, and the subsequent optimisation of interventions.

## Methods

### Participants

Participants were treatment-seeking assault and road traffic accident (RTA) survivors with a primary diagnosis of Posttraumatic Stress Disorder according to DSM-IV-TR (American Psychiatric Association, 2000). Inclusion criteria were exposure to an assault or RTA, and a number of standard fMRI-related safety criteria. They were recruited from an NHS outpatient clinic for anxiety disorders and trauma in London. A trauma-exposed control group without PTSD who had been exposed to an assault or RTA and fulfilled the fMRI safety requirements was recruited via flyers in the community or via recruitment from ongoing research studies at the Institute of Psychiatry, Psychology and Neuroscience, King’s College London, UK. Initial contact was established with 232 individuals, of whom 58 were not interested in taking part in the study, 27 were screened out according to the fMRI safety standards and 58 met other exclusion criteria. Overall, 89 participants were included and three participants dropped out prior to assessment and treatment sessions. From the 86 remaining participants, 10 were excluded due to technical problems with their fMRI data acquisition. Furthermore, two participants were excluded because they did not respond to the therapy. The final sample consisted of 74 participants, of whom 15 were controls, 43 had PTSD at the time of the initial assessment (31 before starting CT-PTSD, and 12 before joining a waitlist), and 16 participants had remitted from PTSD following a course of CT. Of those with PTSD at initial assessment, 14/43 returned for a second scan after a course of CT and 8 returned for the second scan after 3 month wait. The total amount of scans was 96 **(Figure 4a)**. Participants were assessed and scanned individually either before and after a course of trauma-focused cognitive therapy for PTSD (Ehlers et al., 2005), before and after waiting to be treated (waiting list), or just at one time point (trauma-exposed control group without PTSD diagnosis, PTSD patients before or after treatment only). This study design makes possible a mixed longitudinal analysis, including all participants scanned before (n=43) and/or after CT (n=29), as well as a smaller but strictly longitudinal analysis including only participants that attended both visits at the two different time points.

**Figure 4.**
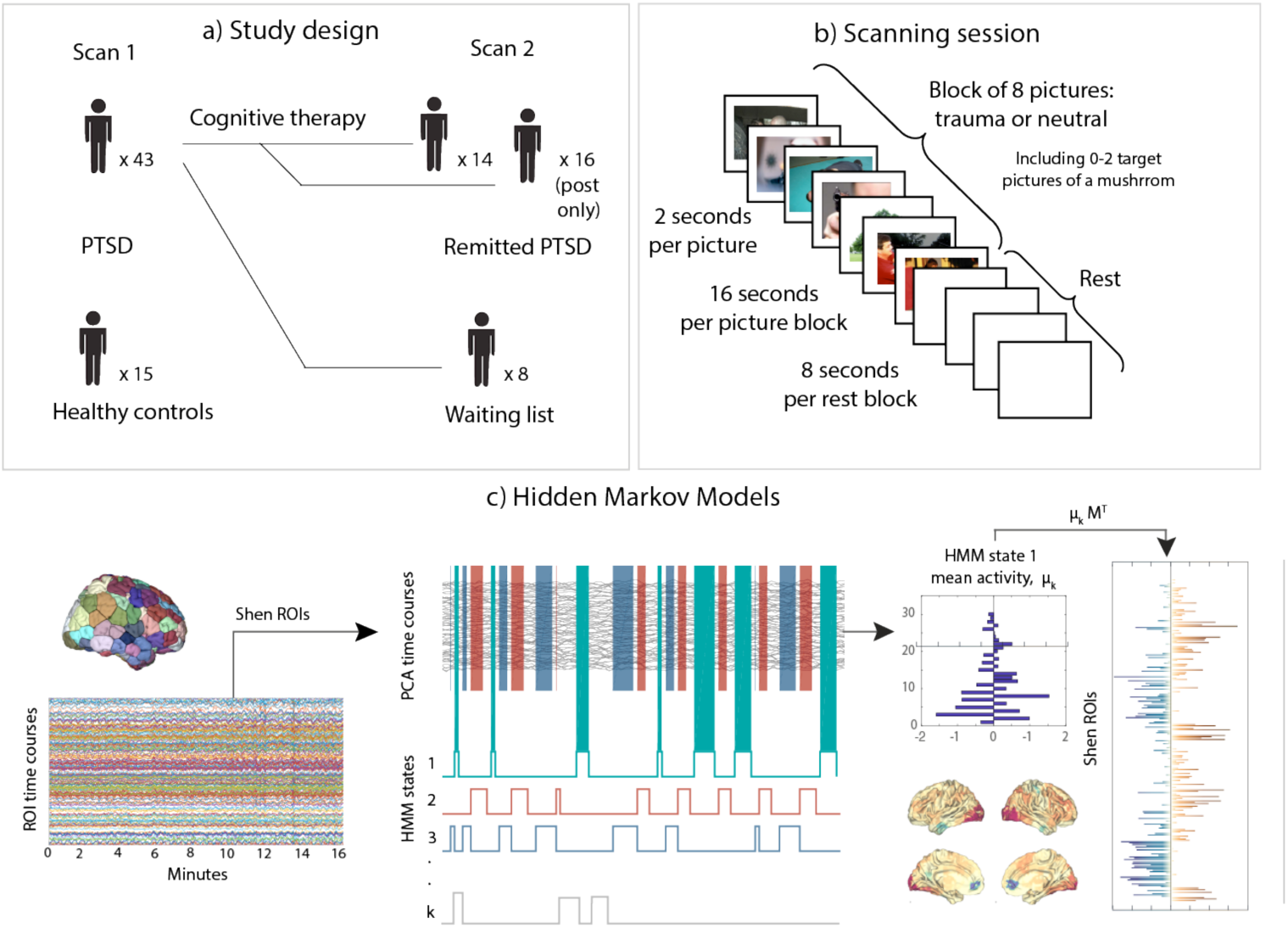
Methods overview. a) Healthy trauma-exposed controls (n=15) and current PTSD participants (n=43) were scanned at baseline. Participants from the PTSD group were scanned a second time after having received cognitive therapy or being on a waiting list condition (n=8) for approximately 3 months (n=31, n=16 only after therapy). b) The task used at both scanning timepoints contained blocks of trauma-related pictures and blocks of neutral pictures. A picture of a mushroom was embedded in each of the blocks 0, 1 or 2 times, to which participants had to react by pressing a button. c) After fMRI data pre-processing, a parcellations with 236 regions (Shen et al., 2013) was applied to extract the timeseries and principal component analysis was performed to reduce dimensionality (75% of variance retained). Subsequently, a Hidden Markov Model was applied, through which we estimated seven different recursive patterns of brain activation and the times at which those patterns dominate brain activity. To visualise each of the patterns, the information was projected back to the Shen space.

Overall, the study included 26 RTA and 48 assault survivors. Average time since the trauma was 4 years and 6.65 months, SD=6.45 years, and the participants mean age was 39.99 years, SD=12.25 years. Some participants were on medication for depression and/or anxiety, see **Supplementary table 1.**

### Procedure

This study was approved by the local ethical review board. Informed consent was obtained from each participant after being given a complete description of the study. Two graduate level psychologists, who were blind to the treatment conditions, conducted semi-structured interviews - the PTSD Symptom Scale Interview (PSS-I, Foa et al., 1993) before and after the treatment or wait list condition. Participants completed the Intrusion Questionnaire for intrusive memories experienced in the scanner (Hackmann et al., 2004) **(Supplementary table 2)**.

### Experimental task

The tasks consisted of 40 blocks of pictures in total, 20 blocks of trauma and 20 blocks of neutral pictures. Order of conditions, i.e., trauma versus neutral, was pseudorandomised. Each picture was presented for 2000ms followed by a 0.4 second blank screen, with an 8 seconds interval between blocks during which a fixation cross was shown, resulting in a total task duration of 16 minutes. Each block consisted of 8 pictures, and within each of these series 0-2 mushroom pictures could appear **(Figure 4b)**. They were asked to press a button box every time they detected a target. However, this information was not used for the analysis.

Stimuli were photographs largely derived from the International Affective Picture System, IAPS (Lang et al., 1997), or from public websites, from various categories, i.e., trauma survivor, injury, trauma-related objects, pre-trauma events (e.g., person sneaking up from behind, a car overtaking in the rain), and pictures depicting actual traumatic events (RTA, assault). Pictures were matched in terms of luminance, number of persons, objects, etc. Parallel categories were introduced for the neutral pictures, i.e., person around the house, household-related objects, two people around the house. Two task versions were constructed, one for assault survivors and one for RTA survivors, with trauma-specific pictures. Pictures across the task had been rated in pilot studies and matched valence on arousal ratings in order to create two versions of each task for the two assessment times (pre- and post-therapy).

### fMRI acquisition and pre-processing

Brain fMRI scanning was performed on a 1.5T Siemens Scanner at the Department of Neuroimaging at King’s College London. Functional images were acquired using an echoplanar protocol (TR/TE/flip angle: 2400/40/80; FOV: 20× 20 cm; matrix size: 96×96) in 36 axial slices (thickness: 3mm; gap: 0.3mm). Data pre-processing was performed using FSL (FMRIB’s Software Library, www.fmrib.ox.ac.uk/fsl) and included head motion correction (MCFLIRT), brain extraction, spatial smoothing with a 5 mm FWHM and highpass filtering (Gaussian-weighted least-squares straight line fitting, with sigma=50.0s). Advanced data denoising was carried out using FIX (Griffanti et al., 2014; Salimi-Khorshidi et al., 2014). Denoised data were first linearly registered to their structural scans and then nonlinearly registered to MNI space. Mean BOLD time-series were then estimated on 268 brain areas of the Shen functional atlas (Shen et al., 2013) by averaging the BOLD signal over all voxels belonging to each brain area **(Figure 4c)**. From these 268 brain regions, 32 were excluded due to signal drop out in orbitofrontal and temporal areas. The excluded timeseries belonged to regions of the orbitofrontal cortex, cerebellum and temporal cortex.

### Hidden Markov Model analysis

Hidden Markov Modelling is an analysis technique to describe multivariate time series data (in this case extracted through the Shen functional atlas) as a sequence of visits to a finite number of networks. Each of these networks has a characteristic signature containing both spatial information (i.e., a specific combination of areas showing different degrees of activation relative to the average and forming a spatial map) and temporal information (i.e., a networks time-course showing when these patterns become active). Mathematically, each state is represented by a multivariate Gaussian distribution, which is described by the mean, and, optionally, the covariance. In our case, we used only the mean to drive the states, so that each state has a specific average activation, and one single full covariance matrix for the entire sample. Consequently, our model takes into account the intrinsic relationship between brain regions, but leaves out any subject-specific variations in the covariance and, therefore, the functional connectivity.

The state sequence is modulated by the probability of transition between all pairs of brain states, which is also estimated within the model; that is, the probability of a given state at a specific timepoint *t* depends on the state that was active at the previous timepoint *(t-1).* Here we applied the HMM on all concatenated scanning sessions, that is including controls at visit 1 and PTSD participants at visit 1 and 2, such that we obtained an estimation of the states at the group level **(Figure 4c)**. This then allowed us to derive subject-specific information of when a state becomes active, which is referred to as the state time-course. Based on the state time-course, we computed the activity time for each state (also referred to as fractional occupancy in related work). These are computed as aggregations of the probabilities of being active at each timepoint for each subject. We derived the activity times independently for each task condition (trauma-related and neutral blocks of picture presentation). The HMM was inferred using the publicly available HMM-MAR toolbox, which provides estimates of the parameters of the state distributions, and the probabilities of each state to be active at each time point (Vidaurre et al., 2017a, 2016).

### Characterisation of states

To make a quantitative assessment of the similarity between our data-driven networks and the anatomy and function of networks characterised in previous literature, we performed automatic decoding of our spatial maps using the Neurosynth database and decoder (Gorgolewski et al., 2015; Yarkoni et al., 2011). Through this approach we were able to compute the correlations between the spatial maps (above average and below average maps separately), and anatomical and cognitive keywords extracted from over 14.000 fMRI studies **(Supplementary figure 1)**. In particular, the 25 terms were retrieved through Neurosynth for each of the above- and below-average activation sub-components separately. From those, up to 10 cognitive terms showing the strongest correlations were selected to represent each of the networks **(Supplementary figure 1)**. The obtained correlation coefficients were moderate in absolute terms, but they were comparable to correlation coefficients obtained in other studies (Gorgolewski et al., 2015; Vidaurre et al., 2017b). Results offered support for our initial neuroanatomy-based labelling of the two DMNs as the mtDMN^+^ and the dmPFC DMN^+^ (for more detail see **Supplementary figure** 1a and **1b**, and **Supplementary figure 2**). The remaining five networks were labelled according to the neuroanatomy of their spatial maps as the Prefrontal^-^, the Salience^+^, the Ventral visual stream^+^, the Auditory^+^ and the Visual/frontal^+^ **(Supplementary figure 1c-1g)**. We then visualised the differences and similarities between the two DMNs, by computing: 1) the area of overlap between the two ‘above-average activation’ areas 2) the remaining of mtDMN+ minus dmPFC DMN+ and 3) the remaining of dmPFC DMN+ minus mtDMN+.

### Statistical analysis

Statistical analyses were run using permutations testing, where we generated surrogate data by permuting the labels of the subjects. First, we looked at differences between the main groups (healthy controls, pre-CT PTSD and post-CT PTSD) in the entire sample, i.e., the pre- and post-CT PTSD groups containing participants that completed both sessions and participants that were only scanned before or after CT-PTSD. Note that as a result, some participants were contributing data at a single session and others at both sessions (before and after CT-PTSD). To assess these differences, we used permutation testing with unpaired t-tests as the base statistic, ignoring that some of the participants were contributing data at both sessions. For an overview of average symptom severity, see **Supplementary table 2**.

Then we investigated differences between PTSD and remitted PTSD only on the subsample of participants who completed both fMRI scans, one before therapy and one after therapy (preCT and postCT). As a control for any changes unrelated to therapy, that might occur over time we also evaluated differences in a different subsample of participants who were scanned at two different points in time but without undergoing any therapy in-between scans (preWAIT and postWAIT). This was done using permutation testing with paired t-tests as the base statistic. Results are shown both after correcting across multiple comparisons, considering both number of states and of experimental conditions, and before correction for multiple comparisons. We used False Discovery Rate (FDR) (Benjamini and Hochberg, 1995) to account for multiple comparisons.

A second statistical analysis was run to test for linear relationships between network activity time and PTSD severity variables. First, we tested for any linear relationships between the activity times of each state and PTSD severity as measured through the three DSM-IV symptom clusters. Similarly, we tested for any linear relationships between the activity times of each state and 1) number of intrusive memories elicited by the fMRI task, and 2) degree to which these intrusive memories present flashback qualities (i.e., vividness, distress and feeling of ‘here & now’). Here, in order to integrate different statistical tests into a single p-value, we made use of the non-parametric combination (NPC) algorithm (Vidaurre et al., 2018c; Winkler et al., 2016); which essentially integrates all hypotheses into a single one, such that we test that at least one alternative hypothesis is true. We integrated the p-values across variables of PTSD symptom severity, an aggregated p-value where the hypothesis is whether at least one of the symptom clusters holds a significant linear relation with the activity time of a specific state. Similarly, when testing for linear relationships between activity time for each state and the flashback qualities of intrusive memories, we integrated the p-values across number of intrusive memories and the three related variables (i.e., vividness, distress and feeling of ‘here& now’) using the same procedure. The NPC algorithm to integrate the individual p-values involved a second-level permutation testing procedure; see Winkler et al, (2016) for further details. P-values were corrected across multiple comparisons using familywise error (FWE) correction (Thomas and Hayasaka, 2003).

## Acknowledgements

The study was funded by the Wellcome Trust [069777]. AE is funded by the Wellcome Trust [069777, 200796], the Oxford Health Biomedical Research Centre and a NIHR Senior Investigator Award. M.L.K. is supported by the European Research Council Consolidator Grant: CAREGIVING (615539), Scars of War Foundation, and Center for Music in the Brain, funded by the Danish National Research Foundation (DNRF117). MWW is funded by the Wellcome Trust (203139/Z/16/Z and 106183/Z/14/Z) and an MRC UK MEG Partnership Grant (MR/K005464/1), and is supported by the National Institute for Health Research (NIHR) Oxford Biomedical Research Centre based at Oxford University Hospitals Trust Oxford University. JJT was supported by the Sigrid Juselius Foundation. The views expressed are those of the authors and not necessarily those of the NHS, the NIHR or the Department of Health.

