## Supplementary information for "Effective psychological treatment for PTSD changes the dynamics of specific large-scale brain networks"

|  | **Mixed sample for group comparison** | | | **Longitudinal sample** | | | |
| --- | --- | --- | --- | --- | --- | --- | --- |
|  | Controls (n=15) | PTSD  (n=43) | Remitted from PTSD  (n=30) | preCT  (n=14) | postCT  (n=14) | preWAIT  (n=8) | postWAIT  (n=8) |
| Age  (mean and SD) | 38.11/12.12 | 37.32/12.48 | 38.23/12.55 | 37.21/14.06 | 37.77/13.98 | 42.26/13.26 | 43.43/13.67 |
| Gender (M/F) | 11/4 | 17/26 | 16/14 | 8/6 |  | 3/5 |  |
| Years since trauma  (mean and SD) | 4.48/6.18 | 4.21/6.87 | 3.35/4.12 | 4.33/4.85 |  | 6.59/9.42 |  |
| Trauma type (RTA/Assault) | 4/11 | 15/28 | 11/19 | 4/10 |  | 3/5 |  |
| Medication (SNRI, SSRI, beta blockers, tricyclic antidepressants) | 1 (6.7%) | 12 (27.9%) | 8 (26.7%) | 6 (42.9%) | 6 (42.9%) | 1 (12.5%) | 1 (12.5%) |

**Supplementary Table 1. Demographic information for all groups.**

|  | **Mixed sample for group comparison** | | | **Longitudinal sample** | | | |
| --- | --- | --- | --- | --- | --- | --- | --- |
|  | Controls (n=15) | PTSD  (n=43) | Remitted from PTSD  (n=30) | preCT  (n=14) | postCT  (n=14) | preWAIT  (n=8) | postWAIT  (n=8) |
| PSSI re-experiencing  (mean and SD) | 0/0 | 11.10/4.02 | 1.23/2.37 | 10.29/3.77 | 1/1.04 | 11.63/4.64 | 8.88/4.91 |
| PSSI avoidance | 0.53/1.19 | 13.01/4.02 | 2.2/2.37 | 11.64/3.30 | 2/2.72 | 13.44/4.17 | 11.63/4.41 |
| PSSI arousal | 0.53/1.41 | 11.16/2.57 | 2.21/2.41 | 11.07/1.44 | 2.29/2.49 | 9.75/4.17 | 11.38/3.58 |
| PSSI total | 1.07/2.49 | 35.33/8.43 | 5.59/11.14 | 33/6.36 | 5.29/5.36 | 34.81/12.01 | 31.88/11.14 |
| Percentage of memories in scanner | 5.67/3.61 | 37.87/28.30 | 25.22/26.34 | 32/27.0 | 7.31/ | 28.75/12.46 | 22.86/13.80 |
| Vividness | 35/35.64 | 58.55/25.25 | 37.83/29.38 | 59.58/28.64 | 26.25 | 58.13/23.29 | 68.57/23.40 |
| Distress | 10/6.32 | 56.84/25.35 | 27.82/28.75 | 57.5/29.58 | 11.25 | 53.75/25.04 | 62.86/29.84 |
| ‘Here /Now’ | 1.67/4.08 | 34.76/25.90 | 12.61/27.17 | 27.92/31.58 | 2.5 | 35/23.90 | 45.71/22.99 |

**Supplementary table 2. PTSD symptomatology**

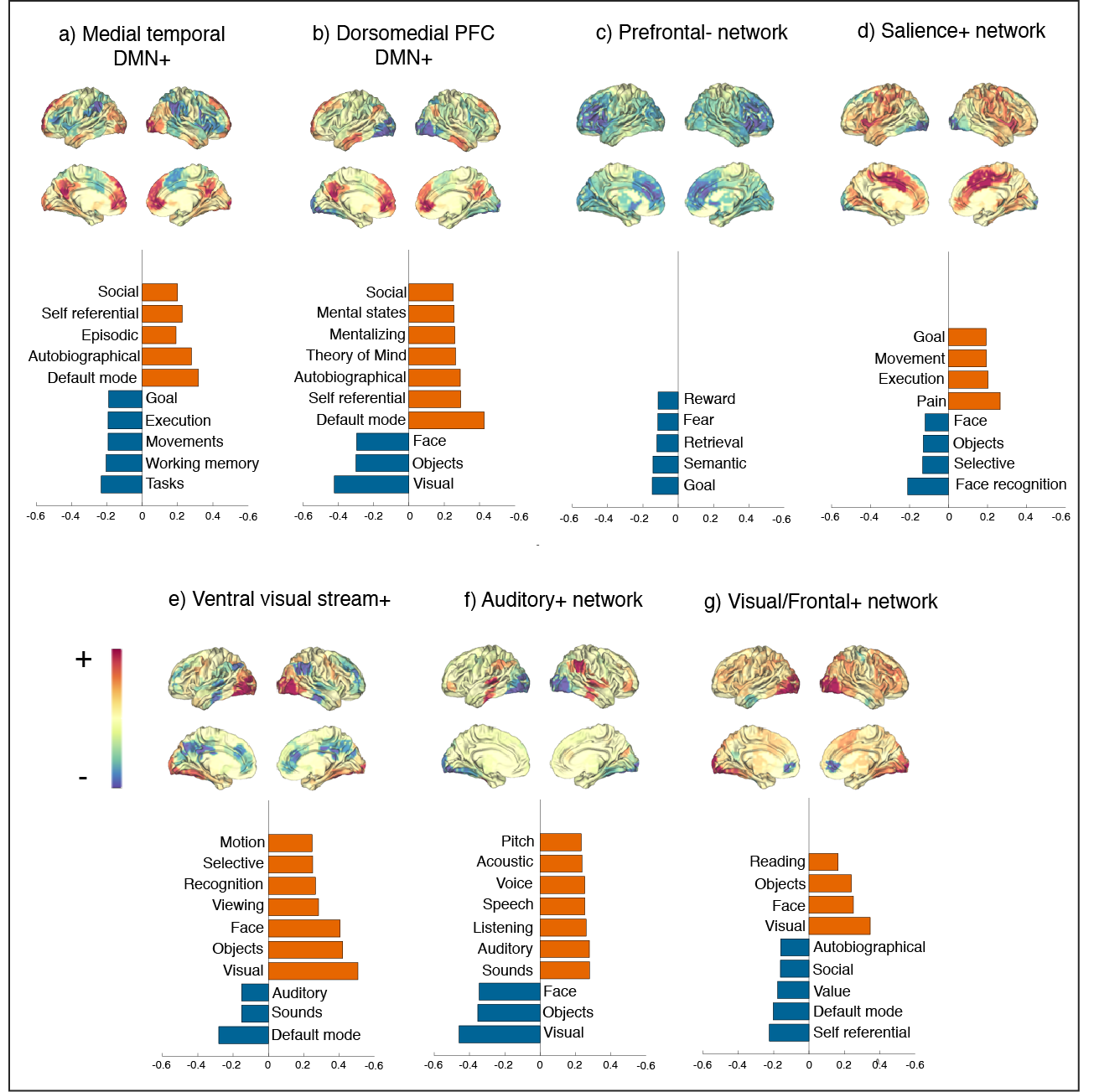

##### **Supplementary figure 1. Correspondence between cognitive terms found in the literature and the areas of above (+/red) and below-average activation (-/blue) for each of the seven networks.** Automatic decoding was performed through Neurosynth (Yarkoni et al., 2011). Up to 10 cognitive terms most strongly correlating with the spatial maps (a-g) are shown below each of the networks.

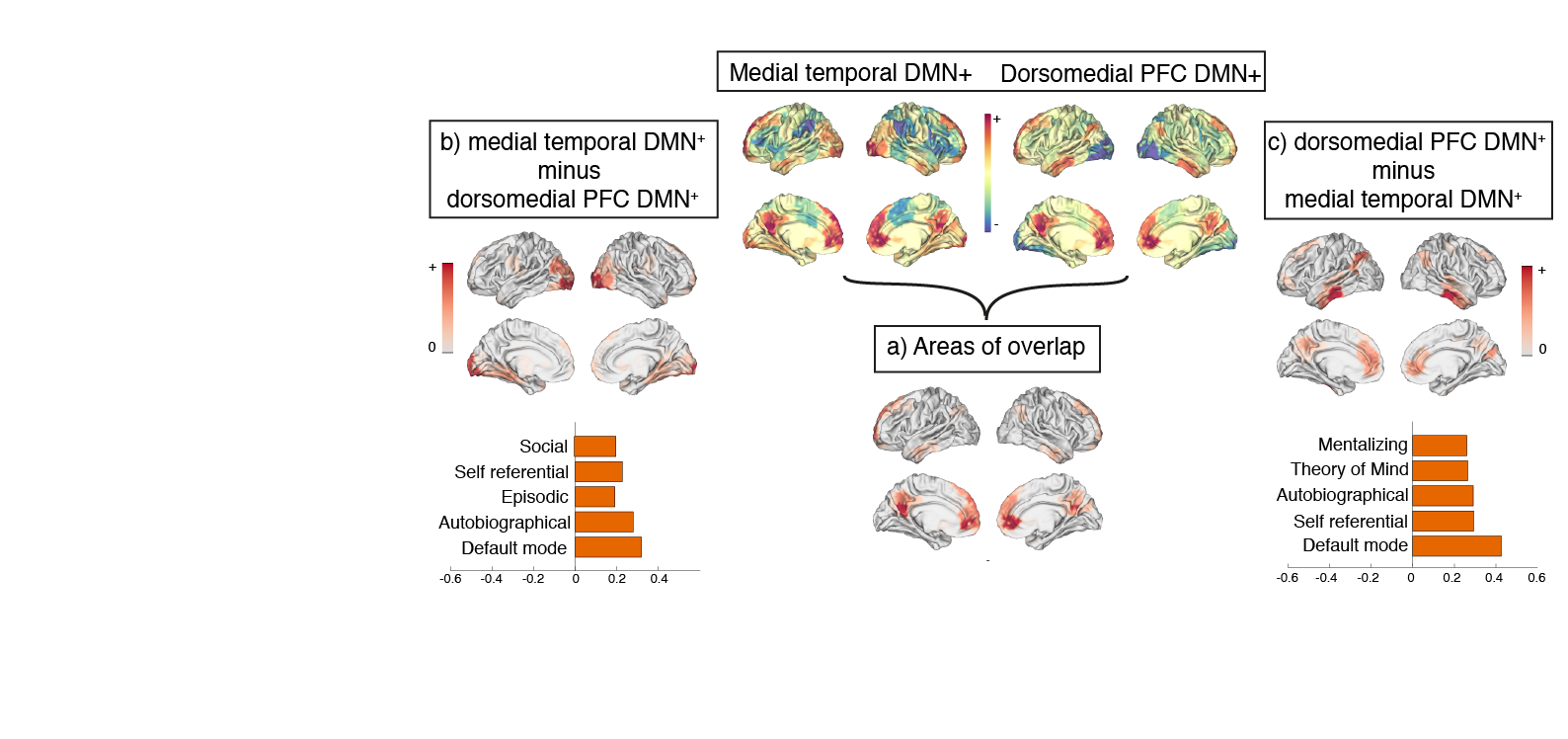

**Supplementary figure 2. Isolation of non-overlapping areas between the two DMNs suggests functional differentiation of their predominant cognitive processes.** a) Areas of overlap involve mainly the anterior medial prefrontal cortex and the posterior cingulate cortex. b) Subtraction of the dmPFC DMN^+^ from the mtDMN^+^ reveals remaining above-average activation in the visual, retrosplenial and parahippocampal cortices as well as in the hippocampal formation. This ‘remaining’ spatial map is most strongly associated with the following terms in the literature: ‘default mode’, ‘autobiographical’, ‘episodic’, ‘self-referential’ and ‘social’. c) Subtraction of the mtDMN^+^ from the dmPFC DMN^+^ reveals remaining above-average activation in the temporoparietal junction, the lateral temporal cortex, anterior and dorsomedial prefrontal cortex and posterior cingulate cortex. This ‘remaining’ spatial map is most strongly associated with the following terms in the literature: ‘default mode’ ‘self-referential’, ‘autobiographical’, ‘theory of mind’, and ‘mentalising’. The mentioned associations are in line with cognitive functions already ascribed to the mtDMN+ (contextualised retrieval of episodic memories) (Cabeza and St Jacques, 2007) and the dmPFC DMN+ (mentalizing and theory of mind) (Mar, 2011); and with another study using the same automatic meta-analysis tool developed by (Gorgolewski et al., 2015; Yarkoni et al., 2011) to relate DMN spatial maps to cognitive functions (Jessica R Andrews-Hanna et al., 2014).

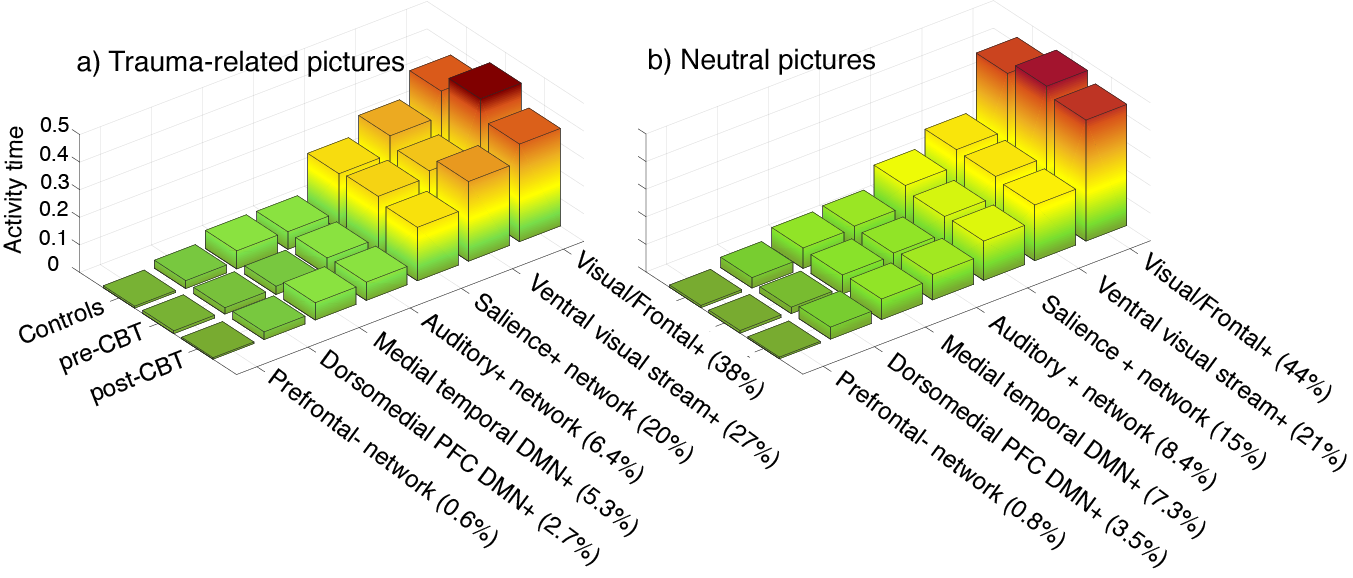

Supplementary figure 3. Healthy controls and patients with PTSD before and after CT spend different amounts of time on each network in the trauma-related and neutral conditions. Activation time for each of the networks is more consistent across groups in neutral (b) than in the trauma-related (a) condition. Furthermore, the smaller percentage of time occupied by activation of the Visual/Frontal^+^ network during presentation of trauma-related pictures (a) in contrast to neutral pictures (b) might be compensated through higher activation times of the salience^+^ network and the ventral visual stream^+^.

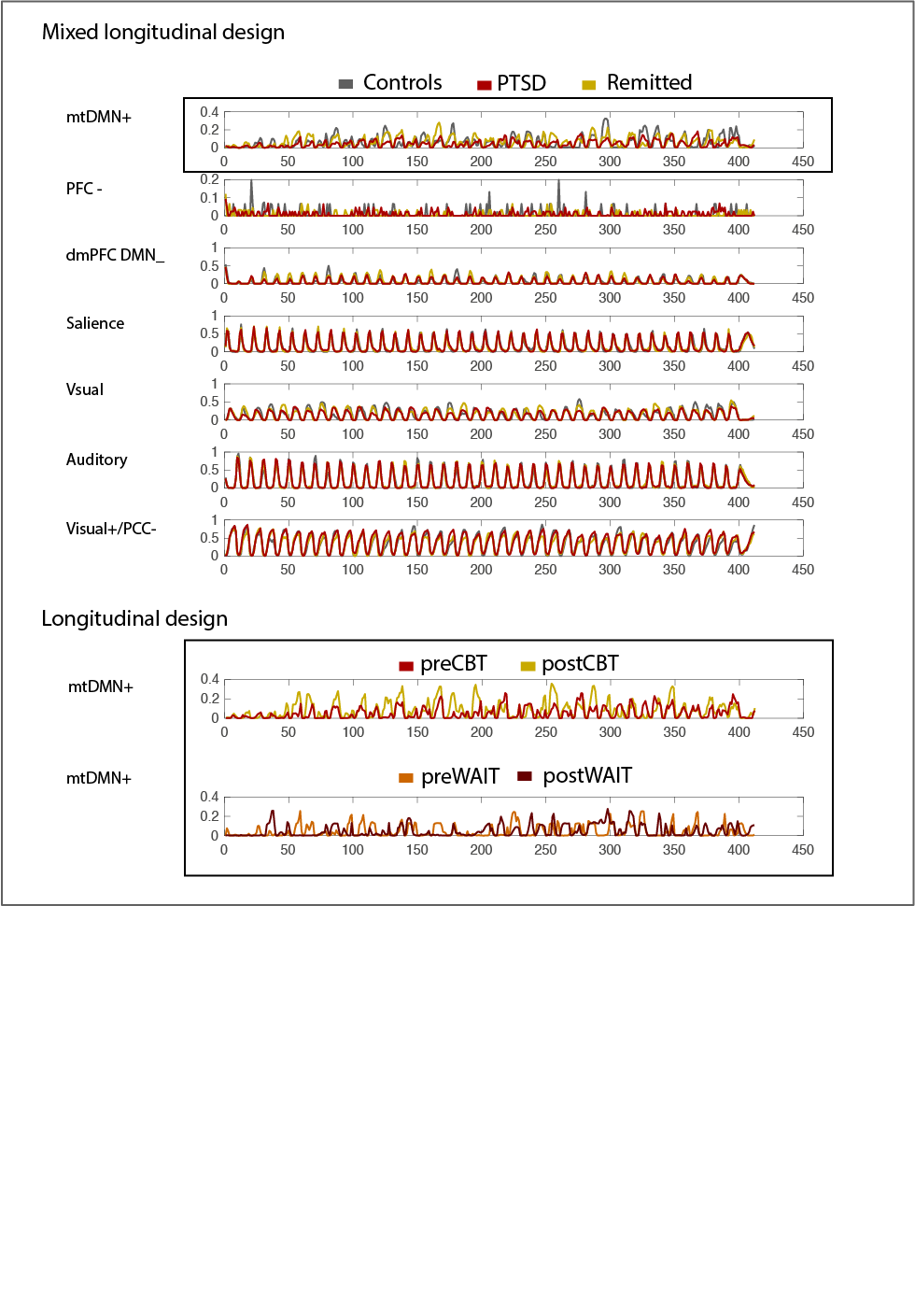

**Supplementary figure 4. Differences in the long-term dependencies of the mtDMN+ might explain the difference observed in activity times of PTSD participants before CT**. Visualisation of the probabilities of activity time for each network and each data acquisition timepoint reveal a trend for increased activity times of the mtDMN^+^ after the first minutes. This trend might be more moderate in pre-CT participants in contrast to healthy controls and post-CT participants (Mixed longitudinal sample, upper panel). The activity times for all the other networks, including the dmPFC DMN^+^, appear to be regular across scanning time and groups. The lower panel shows no signs of a differential trend in pre-WAIT vs. post-WAIT for the mtDMN^+^.
